# Adjunctive inhibition of the integrated stress response pathway accelerates bacterial clearance in a mouse model of tuberculosis

**DOI:** 10.1101/2021.12.31.474642

**Authors:** Stefanie Krug, Pankaj Prasad, Shiqi Xiao, Shichun Lun, Camilo A. Ruiz-Bedoya, Mariah Klunk, Alvaro A. Ordonez, Sanjay K. Jain, Geetha Srikrishna, Igor Kramnik, William R. Bishai

**Author notes:** **Corresponding author**: William R. Bishai. These authors contributed equally (Stefanie Krug, Pankaj Prasad).

## Abstract

Tuberculosis (TB) is a devastating infectious disease that continues to cause millions of human deaths every year. Even though most cases of TB can be cured with a 6-month antibiotic combination therapy, these long treatment durations have led to the emergence of multi-drug resistance and pose a major hurdle to global TB control. Despite numerous advances in TB drug development, a substantially shortened treatment time has yet to be achieved. Given the rise in antibiotic resistance, an alternative strategy to the direct targeting of *M. tuberculosis* (*M*.*tb*) is the development of host-directed therapies (HDTs) that promote bacterial clearance and/or lung health when given adjunctive to standard TB antibiotics. We recently discovered that a small molecule inhibitor of the Integrated Stress Response (ISR), which is abnormally activated in TB and associated with the formation of necrotic granulomas, reduced *M*.*tb* numbers and lung inflammation in mice. Here, we evaluated the therapeutic potential of adjunctive ISR inhibition in the context of standard TB therapy. Throughout the course of treatment, ISR inhibition robustly lowered bacterial burdens compared to standard TB therapy alone and accelerated the time-to-sterility in mice, as demonstrated by significantly reduced relapse rates after 4 months of treatment. In addition, mice receiving adjunctive ISR inhibition tended to have reduced lung necrosis and inflammation. Together, our findings identify the ISR pathway as a promising therapeutic target with the potential of shortening TB treatment durations and improving lung health.

## INTRODUCTION

Despite decades of efforts to understand the disease, and the development of a highly effective first-line treatment regimen, tuberculosis (TB) remains one of the top ten causes of death worldwide and the leading cause of death by an infectious agent. In 2020, about 1.5 million people died of TB, and an estimated 10.4 million were infected by *Mycobacterium tuberculosis* (*M*.*tb*) (1). Among the factors that contribute to the persistently high disease prevalence are the emergence of multi-drug resistance, HIV co-infections, and the lengthy duration of first-line drug therapy (6 months), which hampers accessibility, implementation, and adherence, thereby promoting drug resistance. Therefore, shortening the duration of treatment could significantly improve global TB control. The effort to reduce treatment time may require a better understanding of the disease pathogenesis and the targeting of previously unexplored pathways.

TB is a chronic, often lifelong disease, and *M*.*tb* employs complex strategies to ensure the long-term survival within its human host. A finely tuned concert of pro- and anti-inflammatory responses that is elicited by the host in response to infection contributes to both bacterial containment and persistence within organized host structures called granulomas (2-5). These responses are principal targets for the development of host-directed therapies (HDT) that aim to refocus immune responses toward a more effective elimination of the infection (6-9). The granuloma, while forming an immune microenvironment that contains the infection, also provides a safe niche enabling the prolonged survival and ultimate transmission of *M*.*tb*. TNF is particularly important for both granuloma formation and maintenance and considered essential for TB containment (10). However, several recent studies, including from our group, show that excessive or prolonged TNF stimulation promotes the necrosis of *M*.*tb*-infected, susceptible macrophages through the generation of reactive oxygen species (11, 12), and the proteotoxic stress response and the superinduction of Type I IFN pathways (13), culminating in the aberrant activation of the Integrated Stress Response (ISR). The ISR is a well-established protective host response to viral infections and other stressors aimed at restoring cellular homeostasis by modulating global protein synthesis via the phosphorylation of eukaryotic translation initiation factor 2 alpha (eIF2α) (14, 15); however, excessive or prolonged ISR activation can instead trigger cell death (15). While its role in bacterial infections is only beginning to be appreciated, the ISR transcription factor ATF3 is highly abundant in murine and human TB granulomas, in particular in cells surrounding the necrotic core of the granuloma (16). We recently discovered a causal relationship between type 1 IFN-driven aberrant ISR activation and the susceptibility to TB and necrotic granuloma formation (13). Importantly, we found that ISR inhibition with the small-molecule eIF2α phosphorylation inhibitor ISRIB showed a therapeutic benefit in *M*.*tb*-infected mice, where it reduced granuloma necrosis, lung inflammation and bacterial numbers over the course of 8 weeks (13). In this study, we followed up on these findings and set out to examine the therapeutic potential of ISR inhibition in the context of standard TB chemotherapy. Here, we demonstrate that adjunctive ISRIB, given in combination with traditional first-line anti-TB drugs, significantly reduced both bacterial burdens and relapse rates in mice and may offer additional therapeutic benefit and/or shorten the duration of TB treatment.

## RESULTS

### Adjunctive ISRIB administration reduces lung bacterial burden following M.tb infection

ISRIB, a highly potent ISR inhibitor with an intracellular EC_50_ of 5 nM, exhibits good pharmacokinetic properties and bioavailability (17, 18). It resets proteostasis induced by chronic inflammation in macrophages (13, 19). In *M*.*tb*-infected B6.Sst1^S^ mice, which are known to form necrotic TB granulomas, we found that ISRIB was effective as a monotherapy and reduced bacterial numbers, lung inflammation and granuloma necrosis even though ISRIB had no direct bactericidal activity against *M*.*tb (13)*. Here, we set out to determine if this therapeutic benefit is preserved when ISRIB (0.25 mg/kg by *i*.*p*.) is given in combination with a standard course of the first-line TB antibiotics, isoniazid (H; 10 mg/kg for 6 months), rifampin (R; 10 mg/kg for 6 months) and pyrazinamide (Z; 150 mg/kg for 2 months), in a mouse model of chronic TB (**Figure 1** A). We carried out the study in C3HeB/FeJ mice, since these mice develop heterogenous, cellular and necrotic granulomatous lung lesions following *M*.*tb* infection that recapitulate the range of disease pathology observed in human TB patients more closely than other strains of mice. In the chronic model, infection is allowed to establish for 4-6 weeks prior to the initiation of antibiotic- or host-directed therapy, which more closely resembles the clinical situation and allows us to probe the therapeutic impact of ISRIB on established granulomas. The efficacy of combination therapy with and without ISRIB was determined by evaluating bacterial clearance, time-to-sterility, lung pathology, and inflammation. To assess the treatment-shortening potential of adjunctive ISRIB therapy, separate groups of mice were treated for 4, 5 or 6 months and held off-treatment for an additional 3 months following drug administration before evaluating relapse rates.

**Figure 1:**
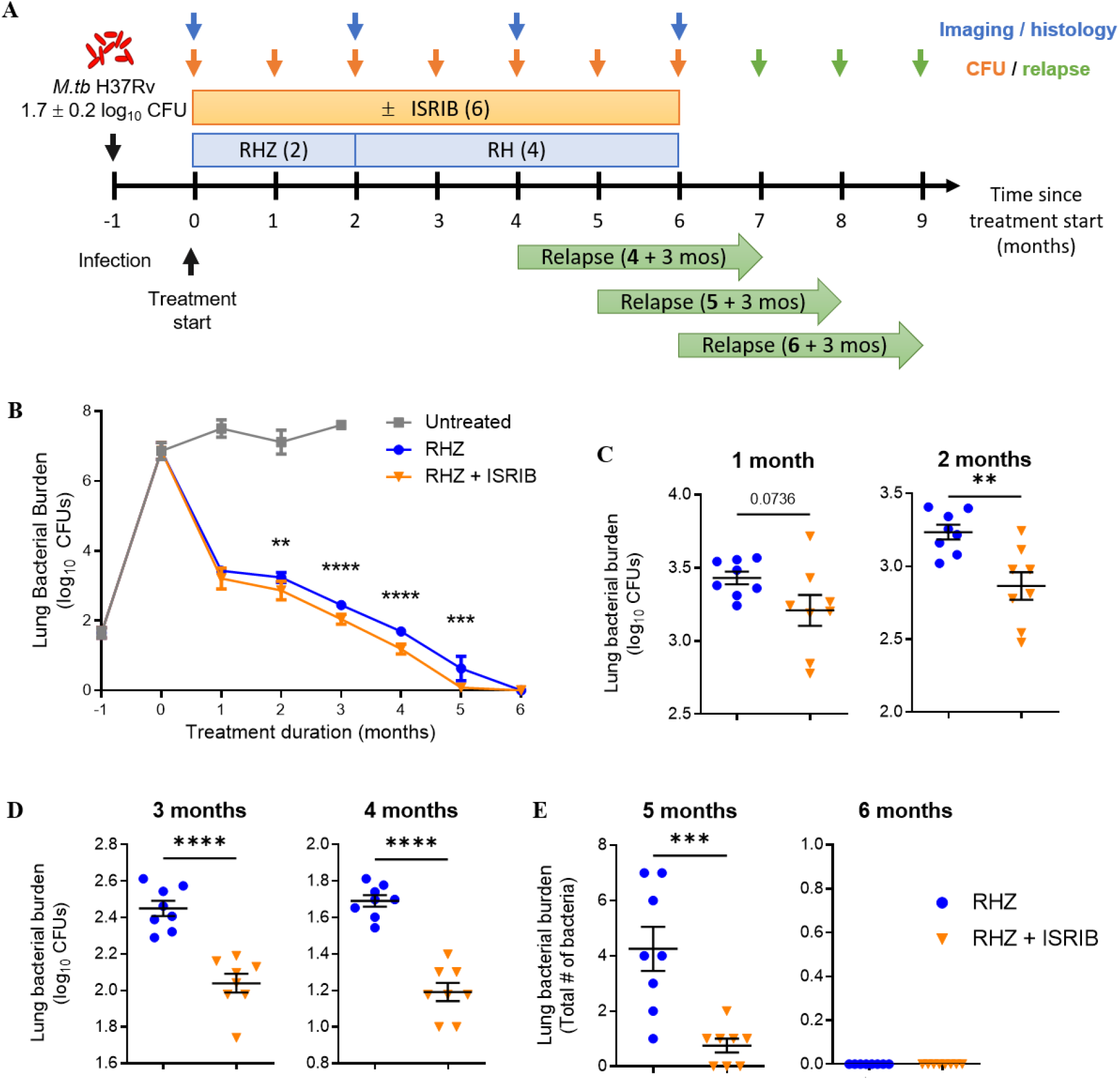
Adjunctive inhibition of the integrated stress response potentiates standard TB therapy. Study overview. Female C3HeB/FeJ mice were aerosol infected with *M*.*tb* H37Rv (implantation: 1.65 ± 0.15 log_10_ CFU). Starting 28 days post infection, mice received standard TB antibiotics (“RHZ”; 10 mg/kg rifampin/R, 10 mg/kg isoniazid/H, 150 mg/kg pyrazinamide/Z for 2 months followed by 4 months of RH) with or without the integrative stress response inhibitor ISRIB (0.25 mg/kg) 5 days/week for up to 6 months. Bacterial burdens were evaluated in 1-month intervals (n=8). After 4, 5 and 6 months of treatment, 15 mice from each group were held without treatment for an additional 3 months to determine relapse rates (green arrows). Lung inflammation and pathology were evaluated at the start of and after 2, 4 and 6 months of treatment by ^18^F-FDG PET/CT and histology, respectively. (**B**) Timecourse of lung bacterial burden (Mean ± SEM og_10_ CFU) in untreated, RHZ-treated, or RHZ + ISRIB-treated mice. (**C-E**) Lung bacterial burden in RHZ or RHZ + ISRIB-treated mice after 1-6 months of treatment. Each symbol represents the log_10_ CFU (**C-D**) or absolute CFU (**E**) of an individual mouse. Lines indicate group mean ± SEM. Statistical differences between RHZ- and RHZ + ISRIB-treated mice were evaluated by unpaired, two-tailed *t*-test. **, *p*<0.01; ***, *p*<0.001; ****, *p*<0.0001. The addition of ISRIB potently reduced bacterial burdens compared to RHZ alone throughout the course of treatment.

At the start of treatment, the mean lung bacterial burden was 6.9 ± 0.2 log_10_ CFU (**Figure 1** B). Infected untreated mice were moribund 4 months after *M*.*tb* challenge and were euthanized. After the first month of treatment, the mean lung CFU declined to 3.4 ± 0.1 log_10_ CFU in mice receiving RHZ, and continued to decline until all mice were culture-negative after 6 months of treatment (**Figure 1** C-E). Even in the context of this highly bactericidal RHZ regimen, the addition of ISRIB robustly enhanced bacterial clearance throughout and significantly lowered lung CFU counts compared to RHZ alone starting from 2 months of treatment. Interestingly, after 5 months of treatment three mice receiving RHZ + ISRIB had no detectable bacilli in their lungs while all RHZ-treated mice had quantifiable bacterial burdens, suggesting that the addition of ISRIB to the first-line regimen may accelerate time to sterility (**Figure 1** E).

### Addition of ISRIB potentiates standard TB therapy and shortens duration of TB treatment

To evaluate the sterilizing effect of adjunctive ISRIB treatment, we also calculated the proportion of mice with culture-positive relapse 3 months after the completion of 4, 5, or 6 months of the indicated drug regimen (see diagram in **Figure 1** A). Mice were considered “cured” if no colonies were found 8 weeks after plating, and “relapsed” if any colonies were found. For mice that received 4 months of the standard regimen (RHZ), there was 100% relapse (15/15) with culture-positive lungs 3 months after the completion of treatment, while relapse was significantly reduced to 33.3% (5/15) in mice that received RHZ + ISRIB (**Figure 2**). After 6 months of treatment, relapse-free cure was observed in all mice that received RHZ + ISRIB while 20% of mice (3/15) that received only the standard regimen still relapsed, indicating that the addition of ISRIB could shorten treatment compared to the control regimen, and more importantly lead to complete sterilization 6 months after treatment.

**Figure 2:**
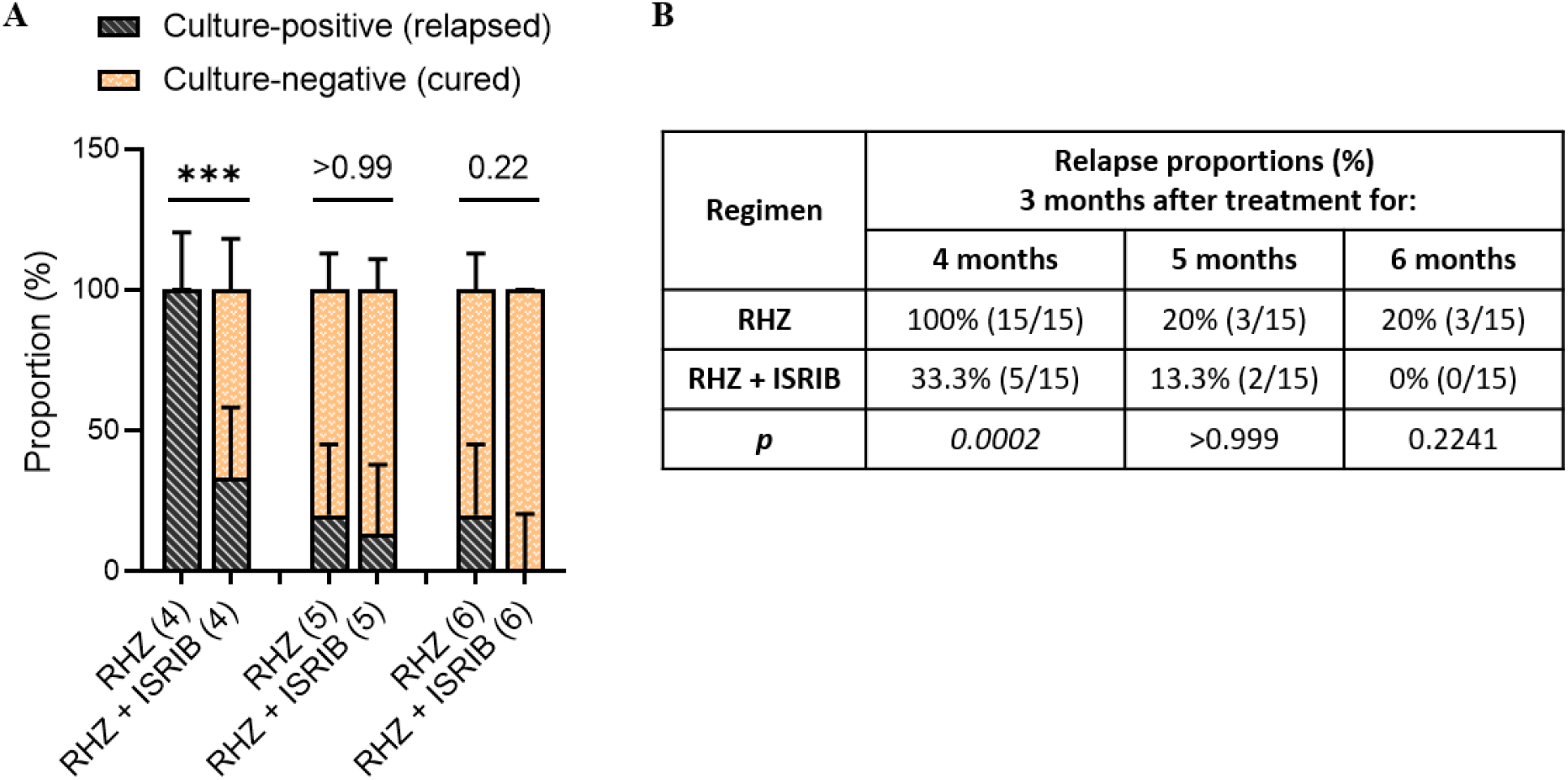
Adjunctive inhibition of the integrated stress response shortens TB treatment duration. Relapse rates in C3HeB/FeJ mice treated with RHZ ± the integrative stress response inhibitor ISRIB (n=15). Mice were treated for 4, 5 or 6 months and held for an additional 3 months without treatments before they were sacrificed and the entire lung homogenized and plated. Mice were considered “cured” if no colonies were detected after incubating plates for 8 weeks, and “relapsed” if any colonies were found. (**A**) Proportion of cured (culture-negative; orange) and relapsed (culture-positive; gray) mice after 2, 4 and 6 months of treatment. Data are presented in stacked columns as fraction of total (%), with error bars indicating 95% confidence intervals computed by hybrid Wilson/Brown method. Relapse proportions were compared by two-sided Fisher’s exact test. (**B**) Table showing relapse percentages, the culture-positive over the total number of mice for each group, and the *p* value calculated for each time point. After 4 months, mice receiving adjunctive ISRIB had significantly lower relapse rates than mice receiving only RHZ.

### Adjunctive inhibition of ISR reduces inflammation and necrosis in mouse lungs

^18^F-fluorodeoxyglucose (^18^F-FDG) accumulates in neutrophils and macrophages at sites of inflammation (20, 21). ^18^F-FDG positron emission tomography co-registered with computed tomography (PET/CT) is therefore extensively used as an imaging modality for evaluating TB lesions in animal models and in patients, and to monitor the bactericidal activity of drugs in preclinical studies (22-25). We therefore evaluated lung inflammation in untreated, RHZ-treated and RHZ + ISRIB-treated mice by serial ^18^F-FDG PET/CT imaging (22). Well-defined foci of ^18^F-FDG uptake colocalizing with TB lesions were observed in lung fields of infected mice, as shown in representative mouse lungs after 2 and 6 months of treatment (**Figure 3** A). Although a time-activity curve of ^18^F-FDG activity calculated from 6 regions of interest (ROIs) per animal did not show statistically significant differences in ^18^F-FDG activity at 2, 4 or 6 months of treatment between RHZ and RHZ + ISRIB-treated mice, the latter group exhibited consistently lower ^18^F-FDG activity than RHZ-only treated mice, suggesting that the addition of ISRIB further reduced inflammation in RHZ-treated mice.

**Figure 3.**
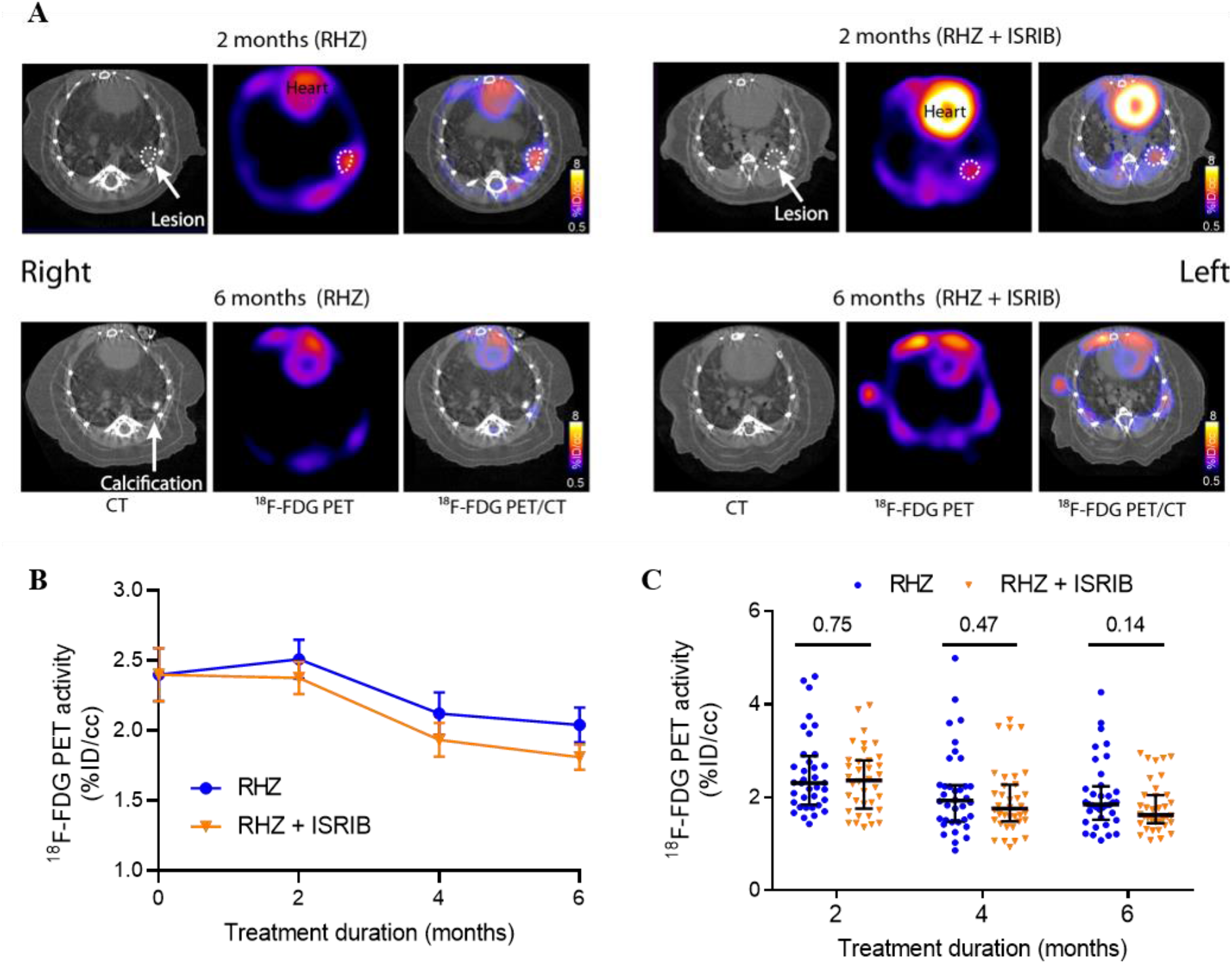
Adjunctive inhibition of the integrated stress response modestly lowers lung inflammation. Lung inflammation in RHZ- and RHZ + ISRIB-treated C3HeB-FeJ mice evaluated by ^18^F-FDG PET/CT (n=6). (**A**) Paired transverse sections (CT, ^18^F-FDG PET and merged ^18^F-FDG PET/CT) in a representative RHZ-treated (left) or RHZ + ISRIB-treated (right) mouse after 2 (top) and 6 (bottom) months of treatment. TB lesions are highlighted with dotted-lines. One mouse from the RHZ group persisted with a calcified lesion. (**B**) ^18^F-FDG PET activity (mean ± SEM) calculated from 6 ROIs per animal. (**C**) Scatter plot of ^18^F-FDG activity per ROI after 2, 4 and 6 months of treatment (6 mice/group, 6 ROIs/mouse). PET data are presented on a linear scale with median ± interquartile range indicated. Differences were assessed by uncorrected Mann-Whitney test. While no differences are noted in the ^18^F-FDG activity at these four time-points, RHZ + ISRIB-treated mice consistently have lower ^18^F-FDG activity than RHZ-only treated mice.

Since a lower ^18^F-FDG activity in the lungs of RHZ + ISRIB-treated mice indicated reduced inflammation, and ISRIB monotherapy led to a significant decrease in necrotic granuloma formation and overall lung destruction 8 weeks post infection as we reported earlier, we performed histological evaluation of inflammation and necrosis in untreated and treated mice. As seen in representative H&E-stained lung sections (**Figure 4;** Supplemental Figures 1-2) of *Mtb*-infected mice treated with RHZ ± ISRIB for 2, 4 or 6 months, or of untreated mice 3 months post infection (control for 2 months treatment), necrotic granulomas were present in the lungs of untreated and RHZ-treated mice but absent in any mice receiving adjunctive ISRIB treatment. These findings suggest that addition of ISRIB to conventional TB therapy may reduce lung inflammation, thereby restoring normal lung structure and function.

**Figure 4:**
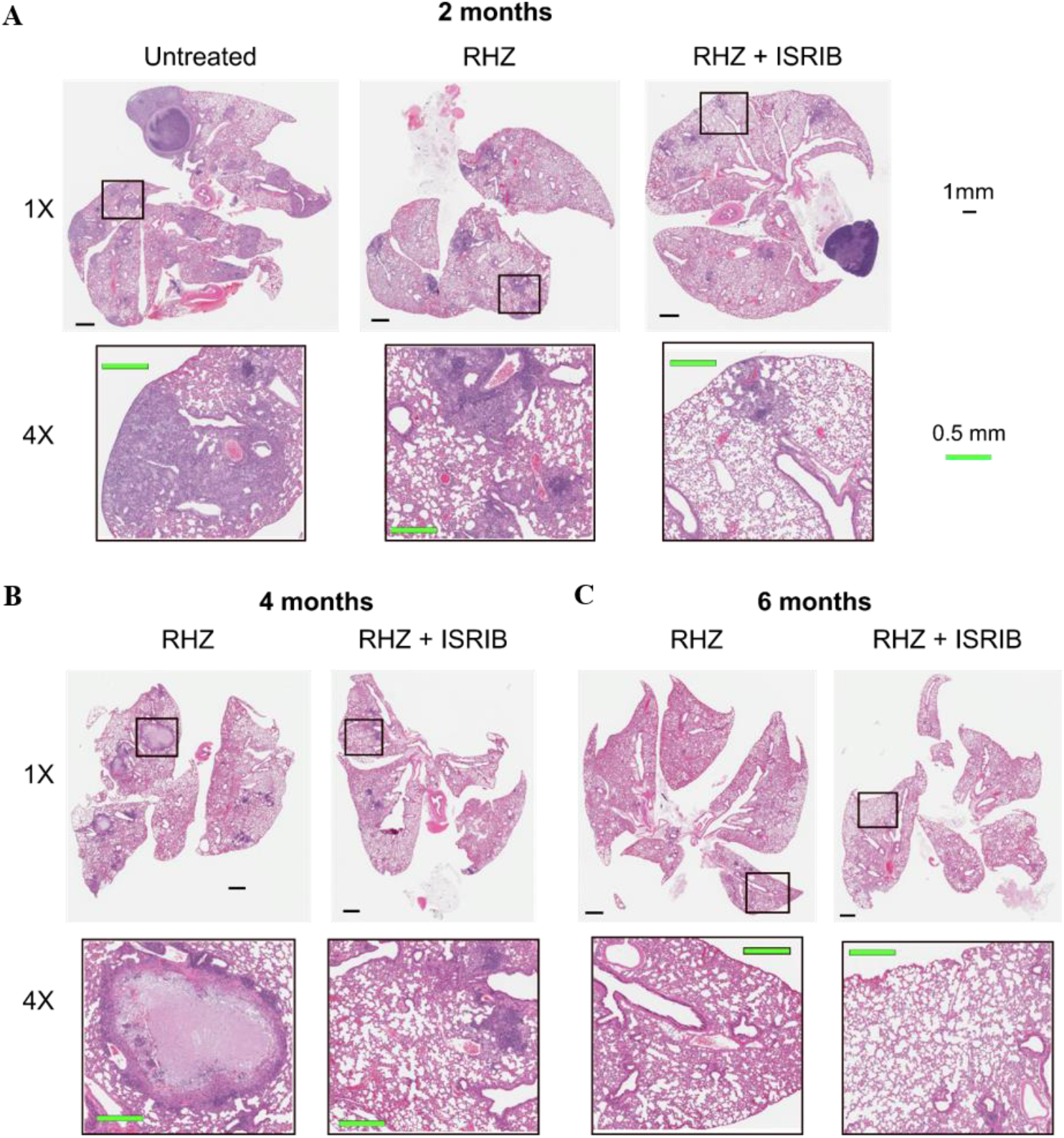
Adjunctive inhibition of the integrative stress response may reduce necrosis in mouse lungs. (**A-C**) Representative H&E-stained lung sections of *M*.*tb*-infected C3HeB/FeJ mice treated with RHZ ± ISRIB for 2 (**A**), 4 (**B**) or 6 months (**C**), or of untreated mice 3 months post infection (control for 2 months treatment) (**A**). The top row of each panel shows low magnification (1X) images of entire lung section. Scale bar indicates 1 mm. The bottom row of each panel shows higher magnification (4X) images of features outlined in the corresponding low magnification image. Scale bar indicates 0.5 mm. Necrotic granulomas were observed in untreated and RHZ-treated, but not in RHZ + ISRIB-treated mice.

## DISCUSSION

The global burden of TB remains unabated and substantial. The existing chemotherapy regimen for TB consists of a 2-month intensive treatment phase consisting of isoniazid, rifampin, pyrazinamide and ethambutol, followed by a 4-month continuation phase with isoniazid and rifampin. Poor adherence to this lengthy treatment regimen has contributed to inefficient disease control and drug resistance, prompting the need for shortened treatment options. Antibiotic resistance can also be circumvented by targeting those host responses to the pathogen that allow for its survival within granulomas.

Granulomas remain a pathological hallmark of TB. Necrotic granulomas, arising from the necrosis of infected macrophages, contribute significantly to disease progression and pathogen spread (26-28). TNF has opposing roles in host-*M*.*tb* interactions. TNF plays a beneficial role in the formation and maintenance of TB granulomas. However, excess TNF promotes necrosis, indicating a pathological role in disease progression. We recently showed that a deleterious macrophage response to prolonged TNF stimulation may drive necrotization of TB granulomas through aberrant, escalating ISR activation driven by the superinduction of the Type 1 IFN pathway, initiated by oxidative stress and dysregulated proteotoxic stress responses, and subsequently amplified by a JNK/IFN-β/PKR circuit (13). Against the backdrop of these dual and opposing effects of TNF, any attempt of antagonizing the necrotic process by directly inhibiting TNF itself might block its protective effects as a mediator of host resistance to infection. We therefore pursued a novel therapeutic strategy of blocking the TNF-induced aberrant stress response using ISRIB, an ISR inhibitor. In fact, our monotherapy studies revealed that ISRIB treatment reduced bacterial loads, inflammation and necrosis in *M*.*tb*-infected mouse lungs compared to untreated mice. This suggested that the addition of ISRIB to the TB regimen could shorten treatment time.

In this study, we therefore tested the efficacy of ISRIB as an adjunctive therapeutic in combination with the standard TB treatment regimen (RHZ). Our studies were carried out in C3HeB/FeJ mice which develop highly organized necrotic lung lesions following *M*.*tb* infection and are therefore particularly valuable in evaluating the efficacy of host-directed TB drugs. We tested the efficacy of ISRIB at a daily dose of 0.25 mg/kg, which provided better inhibition of *M*.*tb* proliferation than higher concentrations of 1.0 mg/kg. At this dose, we found that ISRIB significantly increased bacterial clearance by the first-line anti-TB drugs when tested in combination, starting from 2 months of treatment, suggesting that ISRIB may accelerate time to sterility. Evaluation of relapse after termination of treatment also showed that the addition of ISRIB to standard therapy led to complete sterilization 6 months after treatment. This beneficial effect of ISRIB was accompanied by reductions in inflammation and necrosis in infected lungs compared with the lungs of mice treated with TB antibiotics alone. Our studies suggest that the addition of ISRIB to the standard TB regimen has a therapeutic benefit against *M*.*tb* proliferation and granuloma necrosis.

Although we found a beneficial effect of ISRIB addition to the standard TB treatment in C3HeB/FeJ mice, which develop hypoxic and necrotic lesions with extracellular bacilli, we believe that our results can be extended to other mouse strains such as BALB/c, C3H/HeJ, and C57BL/6 that do not exhibit hypoxic or necrotic lesions, but nevertheless show marked lung inflammation upon infection. In addition, our results are also applicable to infected nonhuman primates which exhibit granulomatous and necrotic lung pathology resembling human TB disease, and hence our results in C3HeB/FeJ mice portend efficacy of ISRIB as adjunctive therapeutic in primate studies and human patients in accelerating bacterial clearance and reducing treatment time.

Activation of the integrated stress response contributes to a wide range of pathologies including neurodegeneration, cancers and infections. Our studies suggest that ISRIB and other inhibitors that are being developed to mitigate ISR in these pathologies may also provide beneficial effects in the form of reduced granuloma necrosis and accelerated bacterial clearance, thereby reducing disease progression and spread in TB.

## ACKNOWLEDGEMENTS

This work was supported by NIH R01 HL133190 and R01 AI153349.

## AUTHOR CONTRIBUTIONS

W.R.B, I.K., S.X. and P.P. conceived the project. W.R.B., P.P., S.K., S.X, S.L., C.A.R-B., S.K.J. and A.A.O. designed the experiments. P.P., S.K., S.X, S.L., C.A.R-B. and M.K. carried out experiments. S.K. analyzed the data and prepared the tables and figures. W.R.B, G.S. and S.K. interpreted the data and wrote the manuscript. All authors read, provided feedback on, and approved the manuscript.

## METHODS

### Mice

Six-week-old female C3HeB/FeJ (stock #658) mice were purchased from The Jackson Laboratory. Mice were maintained under ABSL3 conditions on a 12h:12h light:dark cycle, with rodent chow and water available *ad libitum*. All animal procedures were approved by the Institutional Animal Care and Use Committee of the Johns Hopkins University School of Medicine.

### Bacteria

*M. tuberculosis* strains H37Rv was obtained from the Johns Hopkins Center for Tuberculosis Research, grown to an optical density at 600 nm of approximately 1.0 in Middlebrook 7H9 broth (Gibco) supplemented with 10% (v/v) oleic acid-albumin-dextrose-catalase (OADC; Difco), 0.5% (v/v) glycerol and 0.05% (v/v) Tween 80 (Sigma-Aldrich) and stored in 1 ml aliquots at - 80°C.

### Aerosol infection

Mice were infected via the aerosol route using the Glas-Col Inhalation Exposure System (Terre Haute, IN). A fresh vial of *M. tuberculosis* was used for each infection, thawed on the day of infection and diluted in sterile phosphate-buffered saline (PBS, pH 7.4) at empirically determined factors to achieve the desired inoculum. On the day after infection, 3 mice per cycle were sacrificed to determine the number of CFUs implanted into the lungs. The general appearance and body weight of mice were monitored at least weekly throughout all experiments. All infections, housing of infected mice and handling of infectious materials were carried out under biosafety level 3 containment in dedicated facilities.

### Drug preparation

Rifampin, isoniazid and pyrazinamide were purchased from Sigma-Aldrich. Solutions of rifampin (R; 10 mg/kg), and isoniazid (H; 10 mg/kg) and pyrazinamide (Z; 150 mg/kg) were prepared weekly in distilled water and stored at 4°C. The small molecule inhibitor of eIF2α phosphorylation, ISRIB (0.25 mg/kg) was dissolved in 45% saline, 50% PEG 400 and 5% DMSO (29). All drugs were administered once daily, five days/week, in a total volume of 0.2 ml per treatment. HZ and R were administered by orogastric gavage and ISRIB by intraperitoneal (*i*.*p*.) injection. The HZ solution was gently heated in a 55°C water bath and vortexed to dissolve prior to treating mice, and administered at least 1 h after R to reduce drug interactions.

### Treatment arms and study design

One month after infection, mice were randomly assigned to three treatment arms: standard TB therapy (“RHZ”; 2 months RHZ followed by 4 months RH), standard TB therapy plus adjunctive integrated stress response inhibition (“RHZ + ISRIB”; 2 months RHZ + ISRIB followed by 4 months RH + ISRIB), and untreated (UNT). A subset of mice from each group (UNT: n=5; RHZ ± ISRIB: n=8) were sacrificed at one-month intervals for bacterial enumeration for the first 3 months of treatment, after which point most untreated mice appeared moribund and were euthanized. To evaluate potential treatment-shortening effects of adjunctive ISR inhibition, relapse rates were determined after 4, 5 and 6 months of treatment with RHZ or RHZ + ISRIB (n=15). For relapse studies, mice were sacrificed 3 months after treatment was stopped and the entire lung homogenate from each mouse was plated undiluted on selective 7H11 agar plates (as described below). The presence of colonies on any plate (“culture-positive”) indicated relapse while no detectable growth on any plate after 8 weeks of incubation (“culture-negative”) was considered cured.

### Tissue collection and bacterial enumeration

Mice were sacrificed at predetermined intervals (see diagram in **Figure 1** A). Lungs were aseptically removed, weighed and placed in 2.5 ml sterile PBS for 24-48 h at 4°C, examined for gross pathology and manually homogenized. Homogenates were serial-diluted, and 0.5 ml plated on Middlebrook 7H11 agar (Difco) supplemented with 10% (v/v) OADC, 0.5% (v/v) glycerol, 10 mg/ml cycloheximide, 50 mg/ml carbenicillin, 25 mg/ml polymyxin B and 20 mg/ml trimethoprim (Sigma-Aldrich). Plates were incubated at 37°C for 3-4 weeks before colonies were counted. For relapse studies, plates were checked after 4 and 8 weeks to allow for proliferation of slow-growing bacilli. Colony numbers were adjusted by the plating and dilution factors to estimate the total colony-forming units (CFUs) and are reported as absolute or log-transformed CFUs per lung.

### Histopathology

For histology, intact lungs were fixed by immersion in 10% neutral-buffered formalin for 48 hours, paraffin-embedded, sectioned and stained with hematoxylin and eosin (H&E) stain. Slides were digitally scanned at 40x on an Aperio AT turbo scanner console version 102.0.7.5 (Leica Biosystems). Image files were transferred using Concentriq for Research version 2.2.4 (Proscia Inc) and visualized using Aperio ImageScope version 12.4.0.5043 (Leica Biosystems Pathology Imaging).

### ^18^F-FDG PET/CT Imaging

For ^18^F-FDG PET/CT, six animals from the RHZ or RHZ + ISRIB groups were sequentially imaged at the start of treatment and after 2, 4 and 6 months of treatment, inside in-house developed, sealed biocontainment devices compliant with BSL-3 isolation (22, 25, 30). Mice were fasted for 8 h prior to imaging. Each animal was injected 6.9 ± 0.69 MBq of ^18^F-FDG intravenously via the tail vein. A 15-minute PET acquisition and subsequent CT were performed using the nanoScan PET/CT (Mediso, Arlington, VA). For each animal, six regions of interest (ROIs) were manually selected using CT as a guide and applied to the PET dataset using VivoQuant™ 2020 (Invicro, Boston, MA) for visualization and quantification. Mean lung ^18^F-FDG PET activity was calculated for each mouse (n=6 per treatment group) as the average activity of all VOIs normalized by injected dose (% ID/cc). Data was analyzed non-parametrically and is presented on a linear scale as median and interquartile range (IQ).

### Statistical analysis

Statistical analyses were performed using Prism version 9.2.0 for Windows (GraphPad). Statistical tests used are indicated in the figure legends. CFU counts were log10-transformed prior to analysis up until 4 months of treatment, after which absolute counts were compared. Differences in bacterial burden between RHZ and RHZ + ISRIB groups were assessed by unpaired, two-tailed *t*-test. Relapse proportions were compared by two-sided Fisher’s exact test and are expressed as the fraction of relapsed over the total number of mice per group. 95% confidence intervals for relapse proportions were computed by hybrid Wilson/Brown method. Differences in ^18^F-FDG PET activity were evaluated by nonparametric Mann-Whitney test. A *p* value below 0.05 was considered significant. Data represent mean ± SEM unless otherwise indicated. Histology images were assembled using the open-source vector graphics editor Inkscape for Windows (v. 0.92.4).

## FIGURE LEGENDS

**Figure S1:**
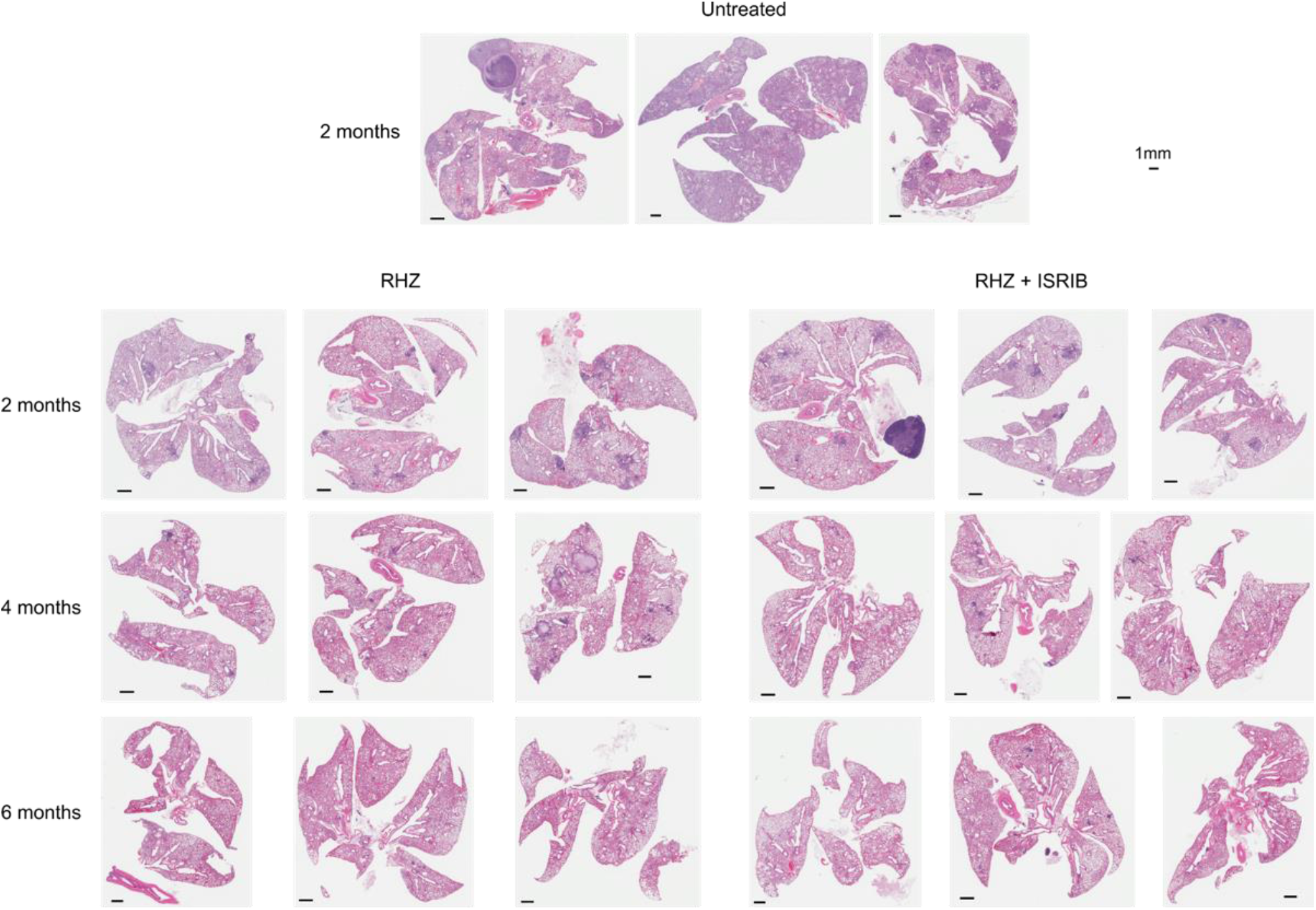
H&E-stained lung sections (low magnification). Low magnification (1X) images of H&E-stained lung sections of *Mtb*-infected C3HeB/FeJ mice treated with RHZ ± ISRIB for 2, 4 or 6 months, or of untreated mice 3 months post infection (control for 2 months treatment). Scale bar indicates 1 mm. Necrotic granulomas were observed in untreated and RHZ-treated, but not in RHZ + ISRIB-treated mice.

**Figure S2:**
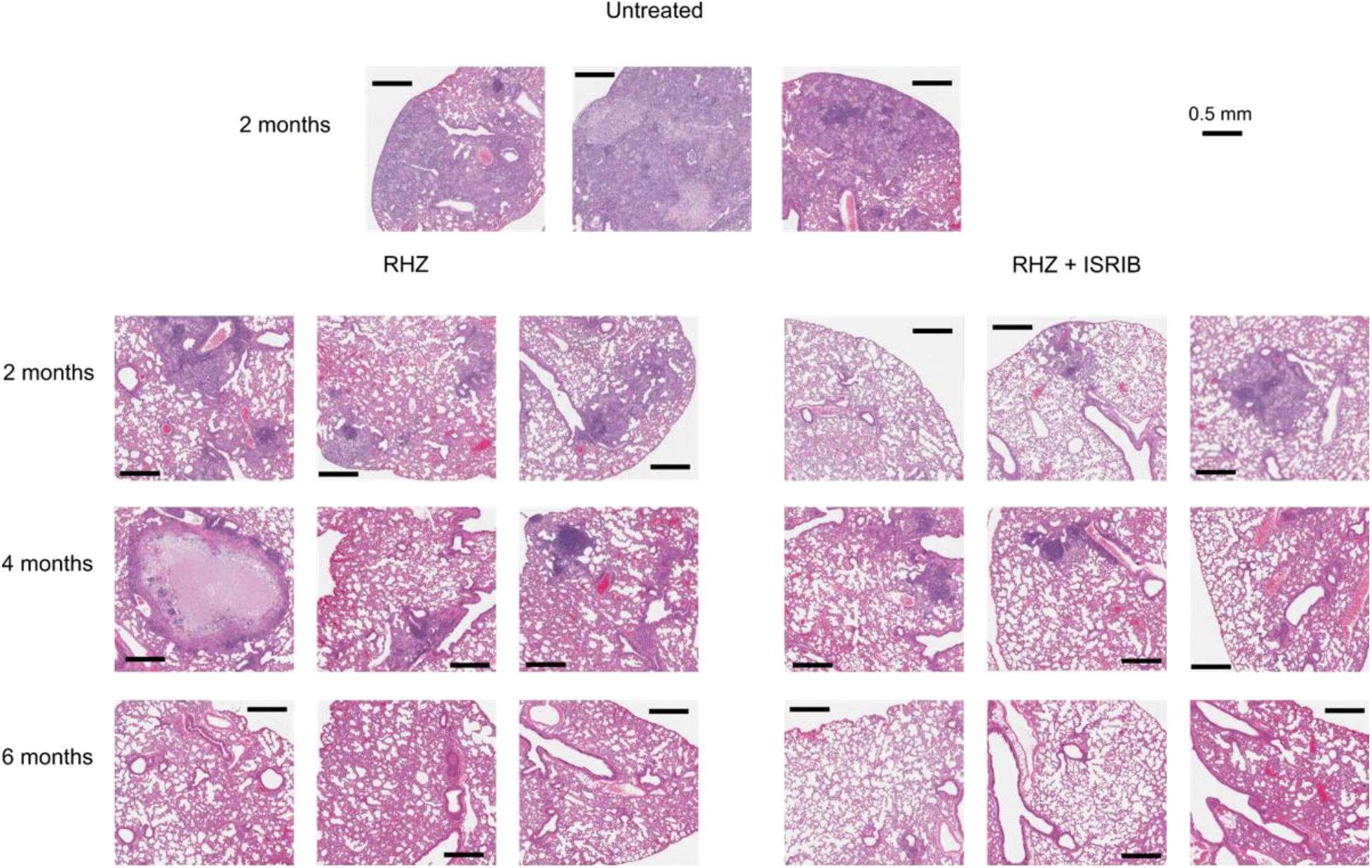
H&E-stained lung sections (high magnification). High magnification (4X) images of H&E-stained lung sections of *Mtb*-infected C3HeB/FeJ mice treated with RHZ ± ISRIB for 2, 4 or 6 months, or of untreated mice 3 months post infection (control for 2 months treatment). Scale bar indicates 0.5 mm. Necrotic granulomas were observed in untreated and RHZ-treated, but not in RHZ + ISRIB-treated mice. The addition of ISRIB may preserve functional airspace in mice.

## Notes

### Competing Interest Statement

The authors have declared no competing interest.

## REFERENCES

1. Global Tuberculosis Report. World Health Organization. 2021.

2. Cambier CJ, Falkow S, Ramakrishnan L. Host evasion and exploitation schemes of Mycobacterium tuberculosis. Cell. 2014;159(7):1497–509. Epub 2014/12/20. doi: 10.1016/j.cell.2014.11.024. PubMed PMID: 25525872.

3. O’Garra A, Redford PS, McNab FW, Bloom CI, Wilkinson RJ, Berry MP. The immune response in tuberculosis. Annu Rev Immunol. 2013;31:475–527. Epub 2013/03/23. doi: 10.1146/annurev-immunol-032712-095939. PubMed PMID: 23516984.

4. Lin PL, Flynn JL. The End of the Binary Era: Revisiting the Spectrum of Tuberculosis. J Immunol. 2018;201(9):2541–8. Epub 2018/10/24. doi: 10.4049/jimmunol.1800993. PubMed PMID: 30348659; PMCID: PMC6217958.

5. de Martino M, Lodi L, Galli L, Chiappini E. Immune Response to Mycobacterium tuberculosis: A Narrative Review. Front Pediatr. 2019;7:350. Epub 2019/09/12. doi: 10.3389/fped.2019.00350. PubMed PMID: 31508399; PMCID: PMC6718705.

6. Abreu R, Giri P, Quinn F. Host-Pathogen Interaction as a Novel Target for Host-Directed Therapies in Tuberculosis. Front Immunol. 2020;11:1553. Epub 2020/08/28. doi: 10.3389/fimmu.2020.01553. PubMed PMID: 32849525; PMCID: PMC7396704.

7. Krug S, Parveen S, Bishai WR. Host-Directed Therapies: Modulating Inflammation to Treat Tuberculosis. Front Immunol. 2021;12:660916. Epub 2021/05/07. doi: 10.3389/fimmu.2021.660916. PubMed PMID: 33953722; PMCID: PMC8089478.

8. Guler R, Ozturk M, Sabeel S, Motaung B, Parihar SP, Thienemann F, Brombacher F. Targeting Molecular Inflammatory Pathways in Granuloma as Host-Directed Therapies for Tuberculosis. Front Immunol. 2021;12:733853. Epub 2021/11/09. doi: 10.3389/fimmu.2021.733853. PubMed PMID: 34745105; PMCID: PMC8563828.

9. Kilinc G, Saris A, Ottenhoff THM, Haks MC. Host-directed therapy to combat mycobacterial infections. Immunol Rev. 2021;301(1):62–83. Epub 2021/02/11. doi: 10.1111/imr.12951. PubMed PMID: 33565103; PMCID: PMC8248113.

10. Wallis RS, Ehlers S. Tumor necrosis factor and granuloma biology: explaining the differential infection risk of etanercept and infliximab. Semin Arthritis Rheum. 2005;34(5 Suppl1):34–8. Epub 2005/04/27. doi: 10.1016/j.semarthrit.2005.01.009. PubMed PMID: 15852254.

11. Roca FJ, Ramakrishnan L. TNF dually mediates resistance and susceptibility to mycobacteria via mitochondrial reactive oxygen species. Cell. 2013;153(3):521–34. Epub 2013/04/16. doi: 10.1016/j.cell.2013.03.022. PubMed PMID: 23582643; PMCID: PMC3790588.

12. Tobin DM, Roca FJ, Oh SF, McFarland R, Vickery TW, Ray JP, Ko DC, Zou Y, Bang ND, Chau TT, Vary JC, Hawn TR, Dunstan SJ, Farrar JJ, Thwaites GE, King MC, Serhan CN, Ramakrishnan L. Host genotype-specific therapies can optimize the inflammatory response to mycobacterial infections. Cell. 2012;148(3):434–46. Epub 2012/02/07. doi: 10.1016/j.cell.2011.12.023. PubMed PMID: 22304914; PMCID: PMC3433720.

13. Bhattacharya B, Xiao S, Chatterjee S, Urbanowski M, Ordonez A, Ihms EA, Agrahari G, Lun S, Berland R, Pichugin A, Gao Y, Connor J, Ivanov AR, Yan BS, Kobzik L, Koo BB, Jain S, Bishai W, Kramnik I. The integrated stress response mediates necrosis in murine Mycobacterium tuberculosis granulomas. J Clin Invest. 2021;131(3). Epub 2020/12/11. doi: 10.1172/JCI130319. PubMed PMID: 33301427; PMCID: PMC7843230.

14. Abdel-Nour M, Carneiro LAM, Downey J, Tsalikis J, Outlioua A, Prescott D, Da Costa LS, Hovingh ES, Farahvash A, Gaudet RG, Molinaro R, van Dalen R, Lau CCY, Azimi FC, Escalante NK, Trotman-Grant A, Lee JE, Gray-Owen SD, Divangahi M, Chen JJ, Philpott DJ, Arnoult D, Girardin SE. The heme-regulated inhibitor is a cytosolic sensor of protein misfolding that controls innate immune signaling. Science. 2019;365(6448). Epub 2019/07/06. doi: 10.1126/science.aaw4144. PubMed PMID: 31273097.

15. Pakos-Zebrucka K, Koryga I, Mnich K, Ljujic M, Samali A, Gorman AM. The integrated stress response. EMBO reports. 2016;17(10):1374–95. Epub 2016/09/16. doi: 10.15252/embr.201642195. PubMed PMID: 27629041; PMCID: PMC5048378.

16. Seimon TA, Kim MJ, Blumenthal A, Koo J, Ehrt S, Wainwright H, Bekker LG, Kaplan G, Nathan C, Tabas I, Russell DG. Induction of ER stress in macrophages of tuberculosis granulomas. PLoS One. 2010;5(9):e12772. Epub 2010/09/22. doi: 10.1371/journal.pone.0012772. PubMed PMID: 20856677; PMCID: PMC2939897.

17. Sidrauski C, Acosta-Alvear D, Khoutorsky A, Vedantham P, Hearn BR, Li H, Gamache K, Gallagher CM, Ang KK, Wilson C, Okreglak V, Ashkenazi A, Hann B, Nader K, Arkin MR, Renslo AR, Sonenberg N, Walter P. Pharmacological brake-release of mRNA translation enhances cognitive memory. Elife. 2013;2:e00498. Epub 2013/06/07. doi: 10.7554/eLife.00498. PubMed PMID: 23741617; PMCID: PMC3667625.

18. Sidrauski C, Tsai JC, Kampmann M, Hearn BR, Vedantham P, Jaishankar P, Sokabe M, Mendez AS, Newton BW, Tang EL, Verschueren E, Johnson JR, Krogan NJ, Fraser CS, Weissman JS, Renslo AR, Walter P. Pharmacological dimerization and activation of the exchange factor eIF2B antagonizes the integrated stress response. Elife. 2015;4:e07314. Epub 2015/04/16. doi: 10.7554/eLife.07314. PubMed PMID: 25875391; PMCID: PMC4426669.

19. Zyryanova AF, Kashiwagi K, Rato C, Harding HP, Crespillo-Casado A, Perera LA, Sakamoto A, Nishimoto M, Yonemochi M, Shirouzu M, Ito T, Ron D. ISRIB Blunts the Integrated Stress Response by Allosterically Antagonising the Inhibitory Effect of Phosphorylated eIF2 on eIF2B. Mol Cell. 2021;81(1):88–103 e6. Epub 2020/11/22. doi: 10.1016/j.molcel.2020.10.031. PubMed PMID: 33220178; PMCID: PMC7837216.

20. Kubota R, Yamada S, Kubota K, Ishiwata K, Tamahashi N, Ido T. Intratumoral distribution of fluorine-18-fluorodeoxyglucose in vivo: high accumulation in macrophages and granulation tissues studied by microautoradiography. J Nucl Med. 1992;33(11):1972–80. Epub 1992/11/01. PubMed PMID: 1432158.

21. Jones HA, Marino PS, Shakur BH, Morrell NW. In vivo assessment of lung inflammatory cell activity in patients with COPD and asthma. Eur Respir J. 2003;21(4):567–73. Epub 2003/05/24. doi: 10.1183/09031936.03.00048502. PubMed PMID: 12762337.

22. Davis SL, Nuermberger EL, Um PK, Vidal C, Jedynak B, Pomper MG, Bishai WR, Jain SK. Noninvasive pulmonary [18F]-2-fluoro-deoxy-D-glucose positron emission tomography correlates with bactericidal activity of tuberculosis drug treatment. Antimicrob Agents Chemother. 2009;53(11):4879–84. Epub 2009/09/10. doi: 10.1128/AAC.00789-09. PubMed PMID: 19738022; PMCID: PMC2772305.

23. Chen DL, Bedient TJ, Kozlowski J, Rosenbluth DB, Isakow W, Ferkol TW, Thomas B, Mintun MA, Schuster DP, Walter MJ. [18F]fluorodeoxyglucose positron emission tomography for lung antiinflammatory response evaluation. Am J Respir Crit Care Med. 2009;180(6):533–9. Epub 2009/07/04. doi: 10.1164/rccm.200904-0501OC. PubMed PMID: 19574441; PMCID: PMC2742744.

24. Xie YL, de Jager VR, Chen RY, Dodd LE, Paripati P, Via LE, Follmann D, Wang J, Lumbard K, Lahouar S, Malherbe ST, Andrews J, Yu X, Goldfeder LC, Cai Y, Arora K, Loxton AG, Vanker N, Duvenhage M, Winter J, Song T, Walzl G, Diacon AH, Barry CE, 3rd. Fourteen-day PET/CT imaging to monitor drug combination activity in treated individuals with tuberculosis. Sci Transl Med. 2021;13(579). Epub 2021/02/05. doi: 10.1126/scitranslmed.abd7618. PubMed PMID: 33536283.

25. Ordonez AA, Wang H, Magombedze G, Ruiz-Bedoya CA, Srivastava S, Chen A, Tucker EW, Urbanowski ME, Pieterse L, Fabian Cardozo E, Lodge MA, Shah MR, Holt DP, Mathews WB, Dannals RF, Gobburu JVS, Peloquin CA, Rowe SP, Gumbo T, Ivaturi VD, Jain SK. Dynamic imaging in patients with tuberculosis reveals heterogeneous drug exposures in pulmonary lesions. Nat Med. 2020;26(4):529–34. Epub 2020/02/19. doi: 10.1038/s41591-020-0770-2. PubMed PMID: 32066976; PMCID: PMC7160048.

26. Batista LAF, Silva KJS, da Costa ESLM, de Moura YF, Zucchi FCR. Tuberculosis: A granulomatous disease mediated by epigenetic factors. Tuberculosis (Edinb). 2020;123:101943. Epub 2020/08/04. doi: 10.1016/j.tube.2020.101943. PubMed PMID: 32741528.

27. Dorhoi A, Kaufmann SH. Pathology and immune reactivity: understanding multidimensionality in pulmonary tuberculosis. Semin Immunopathol. 2016;38:153–66. Epub 2015/10/07. doi: 10.1007/s00281-015-0531-3. PubMed PMID: 26438324.

28. Kramnik I, Beamer G. Mouse models of human TB pathology: roles in the analysis of necrosis and the development of host-directed therapies. Semin Immunopathol. 2016;38(2):221–37. Epub 2015/11/07. doi: 10.1007/s00281-015-0538-9. PubMed PMID: 26542392; PMCID: PMC4779126.

29. Sidrauski C, McGeachy AM, Ingolia NT, Walter P. The small molecule ISRIB reverses the effects of eIF2alpha phosphorylation on translation and stress granule assembly. Elife. 2015;4. Epub 2015/02/27. doi: 10.7554/eLife.05033. PubMed PMID: 25719440; PMCID: PMC4341466.

30. DeMarco VP, Ordonez AA, Klunk M, Prideaux B, Wang H, Zhuo Z, Tonge PJ, Dannals RF, Holt DP, Lee CK, Weinstein EA, Dartois V, Dooley KE, Jain SK. Determination of [11C]rifampin pharmacokinetics within Mycobacterium tuberculosis-infected mice by using dynamic positron emission tomography bioimaging. Antimicrob Agents Chemother. 2015;59(9):5768–74. Epub 2015/07/15. doi: 10.1128/AAC.01146-15. PubMed PMID: 26169396; PMCID: PMC4538528.

